# Sex and Ageing: The Role of Sexual Recombination in Longevity

**DOI:** 10.1101/029082

**Authors:** Phillip R. Smith

## Abstract

I use a set of machines based on the concept of nested rule systems built on the a modified version of the Wolfram elemental cellular automata to investigate the role of recombination in providing resistance to ageing. Class III and class IV machines are observed to respond differently to recombination. Class IV machines show recombinational centring in their neutral networks whereas class III machines respond negatively to recombination. Rule 110 shows a unusual response to recombination. Recombination selects for resistance to recombination, the population moves to regions of genome space with high redundancy, this results in organisms with highly robust genomes, more likely to complete development and to be long lived. The increase in longevity may be sufficient to compensate for the costs of sex, including the two fold cost of sex, through increased reproductive potential in long lived organisms requiring long maturation times. Large complex species should therefore be resistant to invasion by asexual mutants whereas small simple organisms with early maturation should be vulnerable to invasion by asexual forms.

## 2 Introduction

### 2.1 The Two Fold Cost of Sex

Why sex? is an often asked question in biology (1). While sex is ubiquitous, found in plants, from the most primitive to the most advanced and in animals both invertebrate and vertebrate. With fungi the situation in more complex with sexual and asexual forms being common. However in large complex multicellular organisms, sex is dominant, asexual forms are found in vertebrate organisms they are reasonably rare.

Sex should be less favourable due to the two fold cost of sex (2). An asexual mutant that only produced females in a population of sexual organisms should out compete the sexual types. Only the female half of the offspring of a sexual individual are capable producing offspring, where as, all of the offspring of an asexual population can reproduce. A population of sexually reproducing individuals should be vulnerable to invasion by a female mutant that produces only females that reproduce asexually.

There are additional costs of sex, recombinational load where individuals perish because of non-viable combinations of genes, also there are all the additional costs involved finding mates.

There are four main theories trying to explain the evolution of sex

1. The tangled bank. The environment is highly variable and thus many combinations of genes are required to so that some may be fitter in at least some of the environments available. Thus sex is seen as increasing variability in offspring.
2. Novel combinations. Genes that are found in different individuals may be much fitter if found in one organism. It is argued that in asexual populations it is harder for new gene combinations to arise.
3. Red Queen hypothesis. This usually argued that recombination is required to respond to constantly changing selection pressures as parasites evolve with a species. The offspring of each individual are genetically unique and therefore the species is able to respond faster to changes in the biological environment.
4. Deterministic Mutation Hypothesis. Mutations accumulate in asexual species via Mullers ratchet. Recombination removes mutations very efficiently by removing mutations that may individually not be harmful but when combined with other mutations with negative effects are fatal.

The Fisher-Muller advantage of sex (3; 4) combines many of these ideas, essentially two beneficial mutations at different loci can only come to be fixed simultaneously in an asexual population if the one mutation occurs only on a chromosome already carrying the other advantageous mutation. In a recombining population, two chromosomes in the population each containing one or other of the mutations can be recombined to produce a chromosome with both beneficial mutations. Otto and Barton (5) find support for this through simulation showing that in finite populations, linkage disequilibria is generated by genetic drift, genome contamination in the deterministic mutation hypothesis. Recombination can act to bring together the beneficial alleles or remove contamination.

The evolution and maintenance of recombination is one of the processes making up the problem of the evolution of sex and the two fold cost of sex. One of the problems for the evolution of recombination is that it can separate good combinations of genes just as effectively as it can bring them together. This process is called recombinational load. It is the portion of mortality that is explained by new bad combinations of genes. The deterministic mutation hypothesis argues that recombinational load can be constructive by removing mutations from the population more efficiently(6). This requires negative epistasis to operate on the combinations of mutations. It is argued by Otto and Gerstein (7) that negative epistasis produces chromosomes that are much less fit and therefore increase recombinational load. It is argued that this recombinational load prevents a sexual mutant invading a population as the recombinational load would inhibit its spread.

### 2.2 Multicellular Organims

Evolutionary theory, perhaps unjustly, is largely focussed on multicellular organisms. This may reflect the more conspicuous nature of the macroscopic world but is in no small part due the greater challenge of explaining how complex multi-cellular organisms, such as ourselves, could have evolved. It is likely, that it is much easier to develop single celled organisms with few cell types than it is to make complex multicellular organisms with multiple organs and differentiated cell types. However, the models used in population genetics, with few exceptions, treat populations as single cells or more precisely single genomes, growth and development are placed in a black box and ignored (8). It is only in organisms where somatic mutations can be inherited, i.e, organisms without a sequestered germ line, that somatic mutation is usually considered to have a role to play in evolution (9). Some attempts to have been made to address this issue, for example, Orr (10) has suggested that somatic mutation selects for diploidy due to the masking effect of an additional functional copy of the gene within the genome.

Growth and development are inescapable consequences of the evolution of multicellularity. While growth can be seen simply as cell proliferation, development requires a complex interplay between cell proliferation, differentiation and apoptosis. Errors in these systems can have a range of consequences, from little or no effect if mutations are late in development, through incorrect or partial fulfilment of the developmental program, to more severe consequences such as loss of control of cell division leading to cancer.

As organisms increase in size they increase in mass with the cube and surface with the square. Resulting in the requirement for complex mechanisms to bring nutrients to cells and remove wastes. Organogenesis is an inevitable requirement for increased mass and is even seen in angiogenesis in cancer tumours. Increased size requires greater internal complexity and therefore greater requirement for integration of components and vulnerability to the gradual disintegration of components. As size increases with the square of cell divisions, not many cell extra cell divisions are required for increases in size.

Each cell division requires DNA replication and during this process there is a certain probability of mutation. The mutation rate per replication, including point mutations and larger scale mutations is unknown. Most current estimates range from 0.018 for *C. elegans* to 0.16 mutations per genomic replication for human (11). More recent studies of mutation rate per reproduction suggest that these rates are significantly underestimated (12) (13). Metazoan growth is by means of asexual reproduction, so if Muller’s ratchet (14) operates in the soma, in theory, mutations should accumulate down cell lineages until a threshold is reached where the cells are dysfunctional or cancerous.

The relationship between cell division and cancer is well understood. Each division introduces both genetic and epigenetic inheritance of cell state and genome. The risk of cancer in human tissues scales with the number of stem cell divisions (15). Cancer progression from a single cell through to a full blown metastatic tumour requires the population overcoming barriers to tumour development. Tumourigenesis requires the tumour to overcome the intrinsic barriers to tumour growth. These have been suggested to be the hallmarks of cancer (16) namely, Self-sufficiency in growth signals, insensitivity to anti-growth signals, evasion of apoptosis, immortalisation, sustained angiogenesis and tissue invasion and metastasis. Growth and development requires controlled cell growth, apoptosis, and controlled movement of cells. Nearly everything that is required to work properly during development is also required to be broken for tumourigenesis.

### 2.3 Neutral Networks

Neutral networks are networks built up from genetic systems where it is practically possible to enumerate every viable genome. Fitness at the level of the individual is meaningless concept as we cannot estimate this parameter. In neutral networks this problem is addresses by limiting “fitness” to a binary system where there is only viable and non-viable. The networks are constructed by connecting genomes together with their neighbours of hamming distance one, that is, they differ by only one character in the string of characters making up the genome.

Neutral networks have been shown to respond to mutation alone, in such systems the mutational load of each node in the network is the number of actual neighbours divided by the number of possible neighbours. Under mutation pressure a population of randomly selected viable genomes moves quickly towards the centre of the network where an equilibrium state is reached between mutation and selection.

Introducing recombination greatly accentuates the compression of the population into the most connected region of the network. This has been observed in model systems of protein folding to produce more thermodynamically stable configurations than mutation alone can (17). RNA folding is another system which has been used to model the behaviour of neutral networks. However these types of models are looking at individual genes, I wanted to look at genomes. In order to investigate this we need some mechanism to generate genomes. This is not a trivial task as we need to find some rational approach to generate the genomes and some way to systematically describe and analyse these genome generating functions.

Living things essentially have two components, their genome and their cellular machinery. We can therefore consider a simple relation ship between these two entities as a string and a machine. The string represents the encoding of all the machinery to build a machine. The machine reads the string and generates a new string and a new machine Fig. 1a

**Figure 1:**
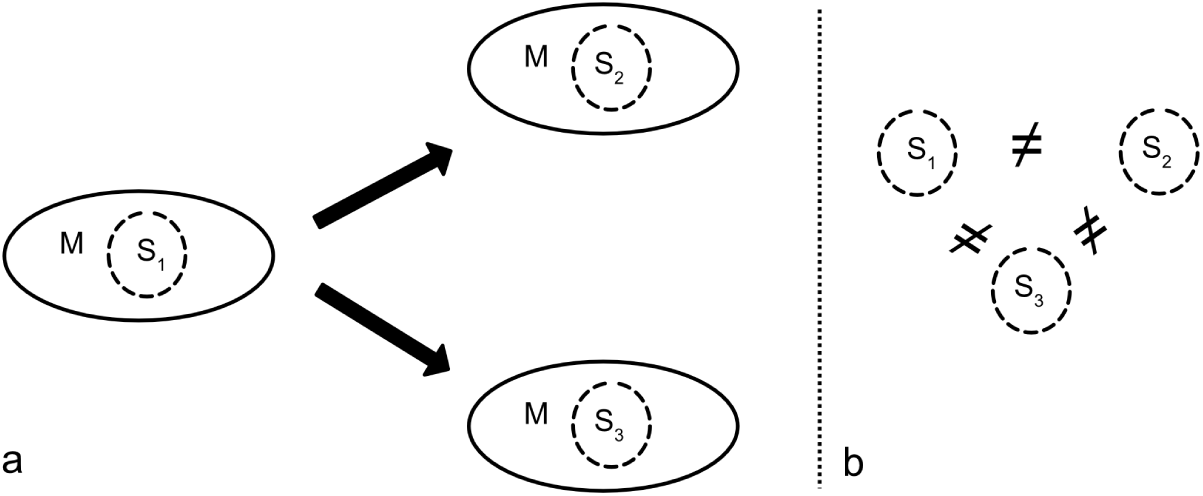
A machine takes a string as an argument and generates a two daughter cells with differing strings to each other and to the parent

If the resulting string and machine is viable we can say that the machine accepts the string. Life can thus be simplified down to a machine that accepts a string that defines a machine. However the process of reproduction is not perfect *s*_1_ ≠ *s*_2_ There is always some difference between the two genomes for any reasonably sized genome Fig. 1b.

If we keep the machine constant and only look at genomes accepted by the machine then for a given genome length we can, in theory, list all viable genomes for that machine. If we connect all these viable genomes together so that all genomes that differ at only one location, hamming distance 1, then the resulting graph is the neutral network (18). This is because all the changes that occurred are neutral they are all equally viable. Genomes are not uniformly distributed in genome space, some regions have a much higher density of solutions than others. Populations of genomes can be bred on this network and the frequency of haplotypes measured. In such networks where there is no difference in fitness between viable individuals there is, however selection for mutational robustness cite (19). As has been shown, elsewhere (20), the frequency of haplotypes is determined by genetic load, which is proportional to density. Those points in the dense regions of genome space have lots of neighbours so mutations, which can be seen as random steps away from a genome, are more likely to find a viable genome. Sparse regions of genome space have very high mutational load, as there is less likelihood of finding a viable neighbour.

### 2.4 Nested Machines

The NK model of Kauffman where there are N genes with K interactions between genes shows that with low values of K fitness landscapes are smooth as K increases, epistasis increases and the fitness landscape becomes more rugged and less correlated (21). Kaufmann, like Wolfram, demonstrated, that there is region between regular and chaotic behaviour where networks of genes are better able to process information. Kauffman used the NK systems to generate truth tables giving fitness values for each combination of genes. This is a similar approach to a neutral network constructed by RNA or protein folding. The NK approach is one way of generating genomes. However I wanted to generate genome sets of variable length using a small finite set of rules and using a method that captured some of the biological structure of living things.

Biological entities are essentially nested machines. Individuals are composed of organs, which are in turn composed of tissues down through cells, organelles, cellular machinery, proteins and other molecules which themselves can be further subdivided. In order to generate enumerable viable genome sets, I built nested machines based on the Wolfram elemental cellular Automata ECA. In effect these machines are simple neural networks with a binary input string of the genome length and a single bit binary output. Classification of the Wolfram ECAs into the four classes is, unfortunately, an undecidable problem (22). Cellular automata can be formally defined using two parameters *N* and *K* unfortunately these are a different N and K to those used by Kauffman in his NK models. Here *N* is the number of neighbours that can be used to decide a cells state and *K* is the alphabet of states a cell can have. In the Wolfram ECAs *N* = 3 and *K* = 2. From these parameters the number of possible neighbour hood states is *K^N^* = 8 The number of possible transition functions or rules is 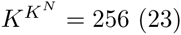.

### 2.5 Cellular Automata

The Wolfram ECA’s use a cylindrical model where the length of the binary string is kept constant. This is done by taking the the first two bits and wrapping around to the last bit to calculate the value of the first bit of the next line. The last bit of the next string is calculated similarly using the last two bits and the first bit. In contrast we want the length of each new line to reduce down eventually down to a single bit. For this reason we did not use the cylindrical model but a simple compression model where each line is two bits smaller than the last line. It is therefore necessary that the strings are all odd numbered and at least three or more bits long. Each layer of the machine is decided by the rule applied to the results of the nested machines above it Fig. 2

**Figure 2:**
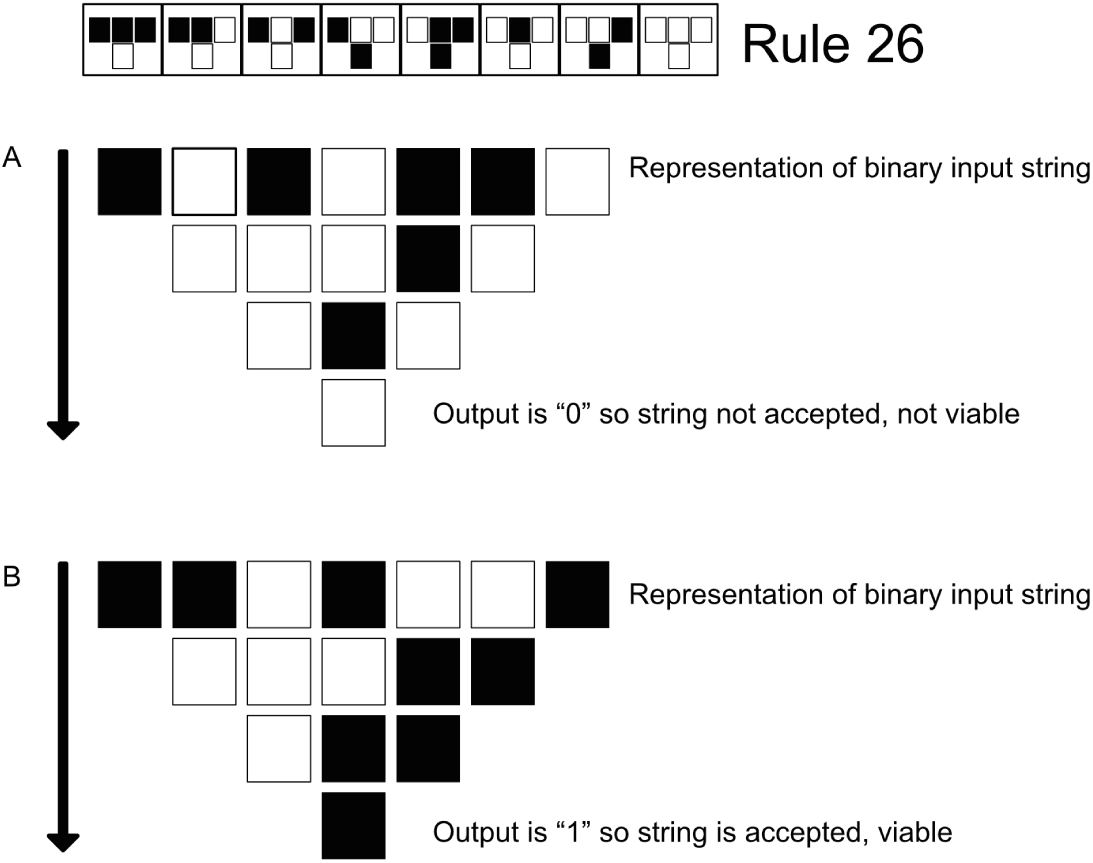
Example A the input string is condensed down to a single bit. The last bit is zero so the string is rejected. In Example B the string is accepted. Both strings were processed using rule 26

This method does have the draw back of edge effects so the behaviour of the 256 rules of the ECA’s may behave differently than they do under the cylindrical model.

There are many advantages in using the Wolfram ECA is that they are well described and researched. There is a finite list of machines that can be systematically investigated. The ECA are not very demanding from a computational point of view so large numbers of trials can be done with varying length genomes and vary population sizes etc.

Wolfram ECA have been suggested to fall in the following four classes

Class I evolves to a homogeneous state.
Class II evolves to simple separated periodic structures.
Class III yields chaotic aperiodic patterns.
Class IV yields complex patterns of localised structures.

## 3 Materials and Methods

Each rule was evaluated by passing all the strings to it of a given length. A string that resulted in the last bit in the process being a 1 was accepted, if it was a 0 then the string was rejected. For a given genome length a one bit heat map can be drawn for every combination of every genome and every rule. Fig. 3 shows all the possible genomes from 00000 to 11111 being evaluated for every possible rule (machine). The rules on the right side of the graph show increasing propensity to accept genomes with the opposite pattern of the left hand side. In the middle, the rules become more complex.

**Figure 3:**
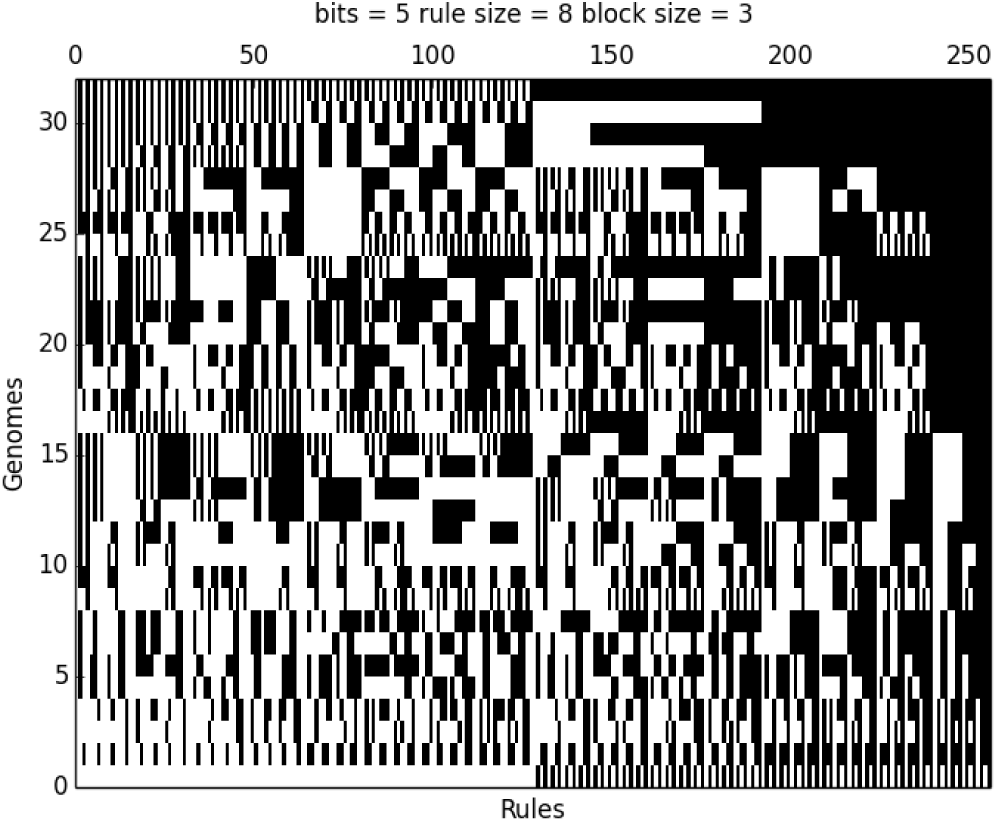
5 bit heat map of every genome for every rule

As can be seen from Fig. 4 increasing the genome size affects the heat map, however the overall pattern remains the same with the lighter patches on the left and darker regions of the right. Some rules at do not accept any string [0,8,32,64,128] whereas some rules like 255 accept every string. The set of accepted strings can be now used to create a neutral network graph. Graphs were generated using the Neato (24) algorithm as part of the graphviz package. Graphs were generated for all the the rules with a genome length of 13. The number of neighbours or the degree of each node was measured and the frequency distribution of node was plotted. The correlation of the degree of a node with its neighbours was measured using a genome length of 21bits. This was done by choosing two connected nodes at random and counting the number of neighbours for each of the two nodes. This graph will be referred to as the NofN graph or correlation. The slope of this graph and the standard error of the correlation was also recorded.

**Figure 4:**
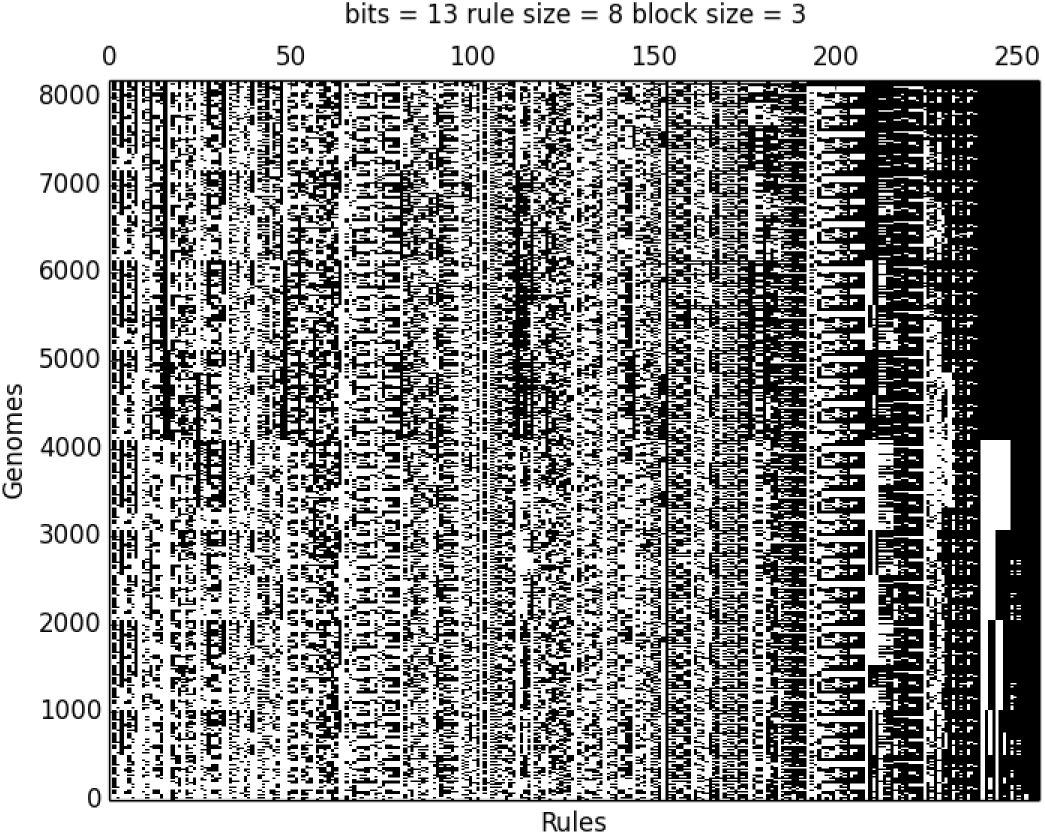
13 bit heat map of every genome for every rule

In addition to the genome sets generated by the rules a set was generated using the sha256 cryptographic hash function. All possible 21 bit genomes were used as arguments for sha256 if the resulting integer from this function was odd then the genome survived if not the genome died. The sha256 algorithm was also used to generate frequency distributions of neighbours and NofN correlation statistics

### 3.1 Breeding experiments

Each rule was then subjected to breeding experiments with various genome lengths. The mutation rate was such that each generation every genome had 1 mutation. Where recombination was included the recombination event occurred after every bit. Lower rates of recombination simply delayed the effect of recombination and took longer to reach equilibrium. The population size was kept at 5000 and period of sexual and asexual reproduction were encountered over the generations. Each experiment was initiated with a random start population of 5000 viable genomes, which underwent sixty generations of asexual reproduction followed by 80 generations of sexual reproduction then a return to asexual reproduction for another eighty generations.

During sexual reproduction two parents were selected at random and both were mutated, a child genome was generated by recombining the two mutant parental genomes. If this was viable the child was added to the next generation. This process was repeated till 5000 individuals were generate for the next generation. The number of unsuccessful trials over the total number of trials was used to calculate the recombinational load of the population which equals 1 − *viability*.

Not all rules reached equilibrium in the time allocated some of the rules were repeated with longer periods of recombination. Effect of genome length was also analysed for some rules and changes in behaviour were noted.

### 3.2 Measuring Resistance to Ageing

For each rule the whole population was sampled at various times during the breeding program. The start population, the population after asexual reproduction, after asexual reproduction and at the end of the run. Each sample was analysed for resistance to ageing by taking every genome in the sample in turn and subjecting it testing.

A population of cells can be simulated in two ways. A liquid culture where a cell lineage can over grow the population and compensate for slow growing lineages. Solid culture where overgrowth of cell line is prohibited. In most multicellular organisms cells are not free to move around and compensate for hypoplasia in distant tissues.

A population of 50 cells in a solid medium was simulated, each cell was mutated if the mutant cell was not viable it was removed from the population Fig. 5. This process was continued till the population reached zero. The number of rounds of mutation taken to kill the population was take as the time of death of the individual. This parameter indicated expected life span of multicellular solid group of 50 cells.

**Figure 5:**
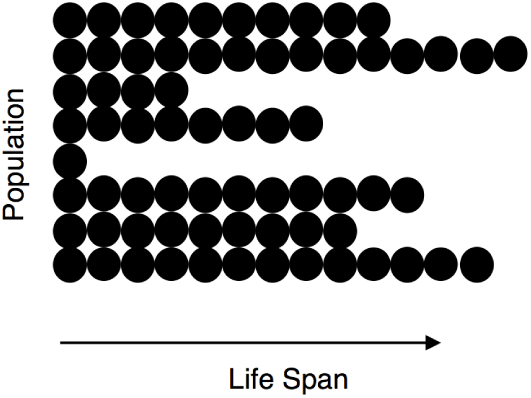
Solid model of cell growth in a multicellular organism

## 4 Results

The cryptographic hash function sha 256 produced a neutral network that was a random graph. The frequency distribution of the number of neighbours was near normal in appearance. The slope of the NofN graph was near zero and the SE was large Fig. 6. Interestingly a slight decease in load under asexual reproduction was observed Fig. 7, that was removed by recombination with load returning to the expected value of 0.5. Suggesting that there are some small clusters of well connected nodes however they are not well connected to each other. Recombination breaks up the genomes of these well connected nodes returning the load to the 0.5 level. In a machine as complex as a pseudo-random number generator (PRNG) that generates deterministic randomness, recombination has a negative affect and asexual reproduction would be favoured.

**Figure 6:**
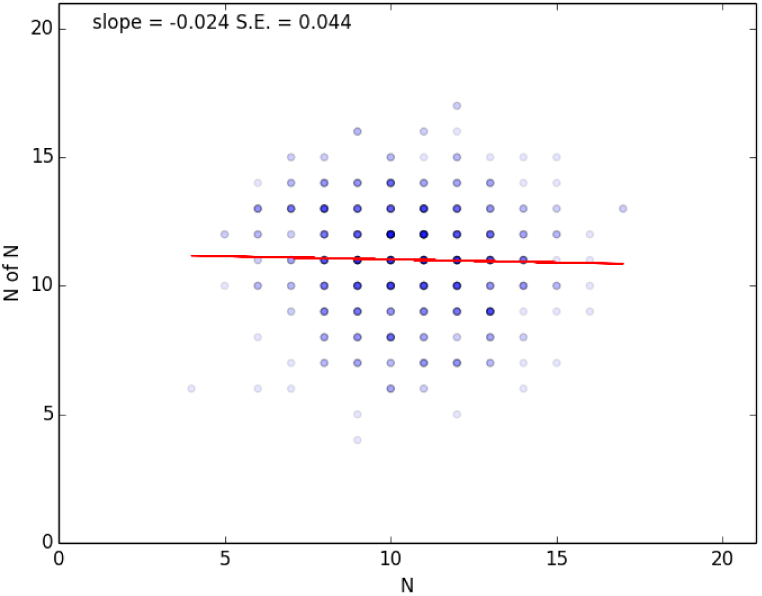
NofN graph for sha256 with 21 bits

**Figure 7:**
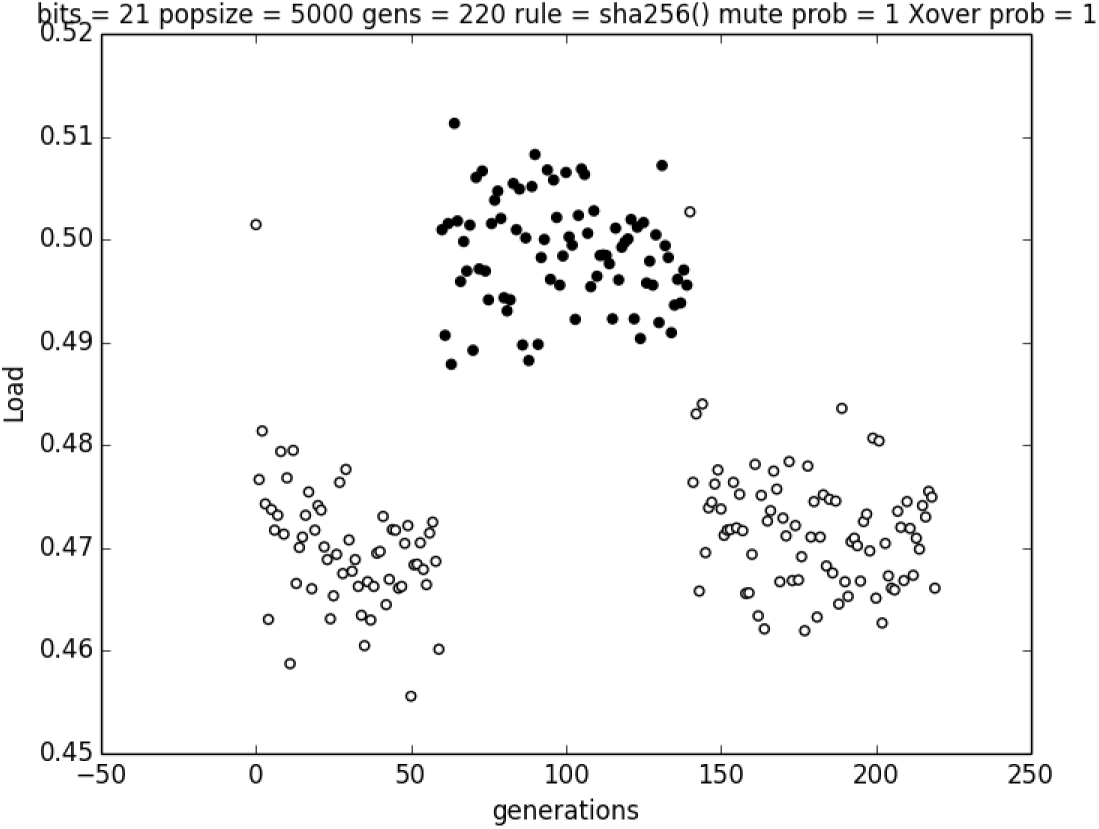
sha256 showed a weak response to mutational load but recombination was unable to reduce recombinational load and increased mutational load

### 4.1 Response to Selection for Mutational and Recombinational load

The nested rule systems for each rule were subjected to sixty generations of asexual reproduction followed by eighty rounds of sexual reproduction after which a return asexual reproduction was performed for a further eighty generations, with the exception of the rules that produced regular graphs of the same degree, all rules responded to mutation by moving towards reduced mutational load. Most rules responded well to recombination Fig. 8

**Figure 8:**
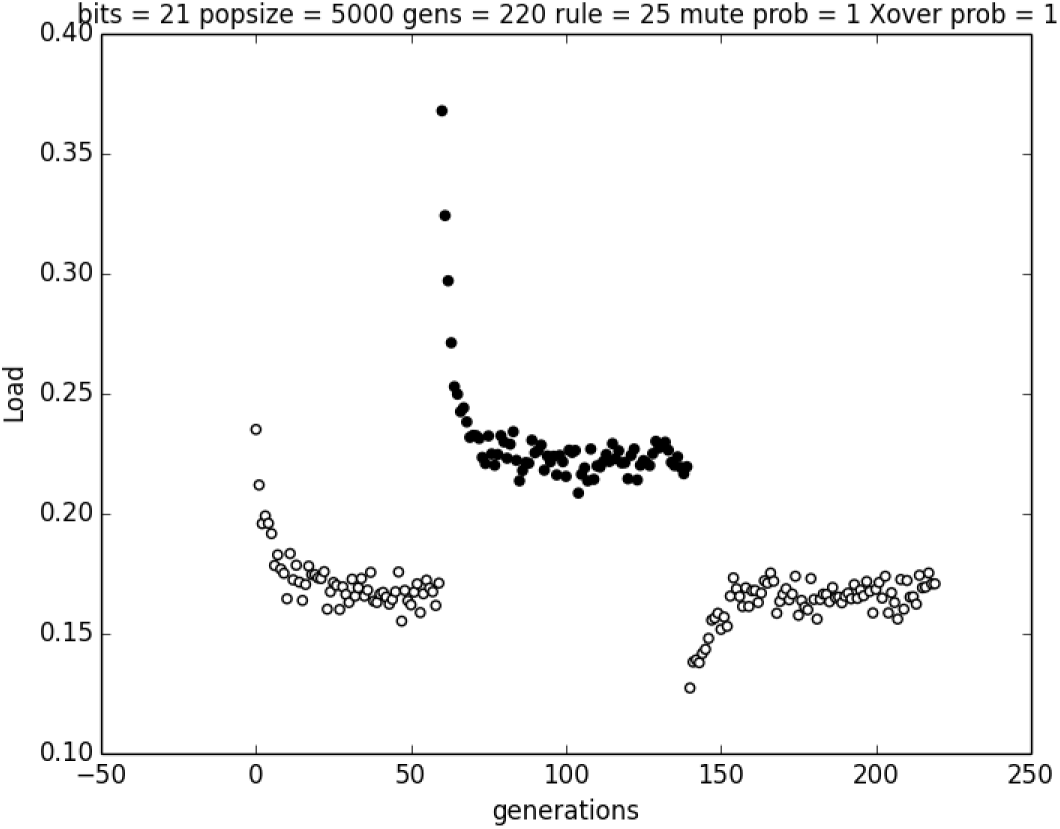
Rule 25 at 21 bits. This rule responds positively with a drop in mutational load following removal of recombination

The reduction in mutational load observed in the rules varied from rule to rule with some having a greater drop in mutational load and others taking longer for that decrease in load to return to the pre-recombination level. The increase in mutation load following recombination was seen in relatively few rules Fig. 9.

**Figure 9:**
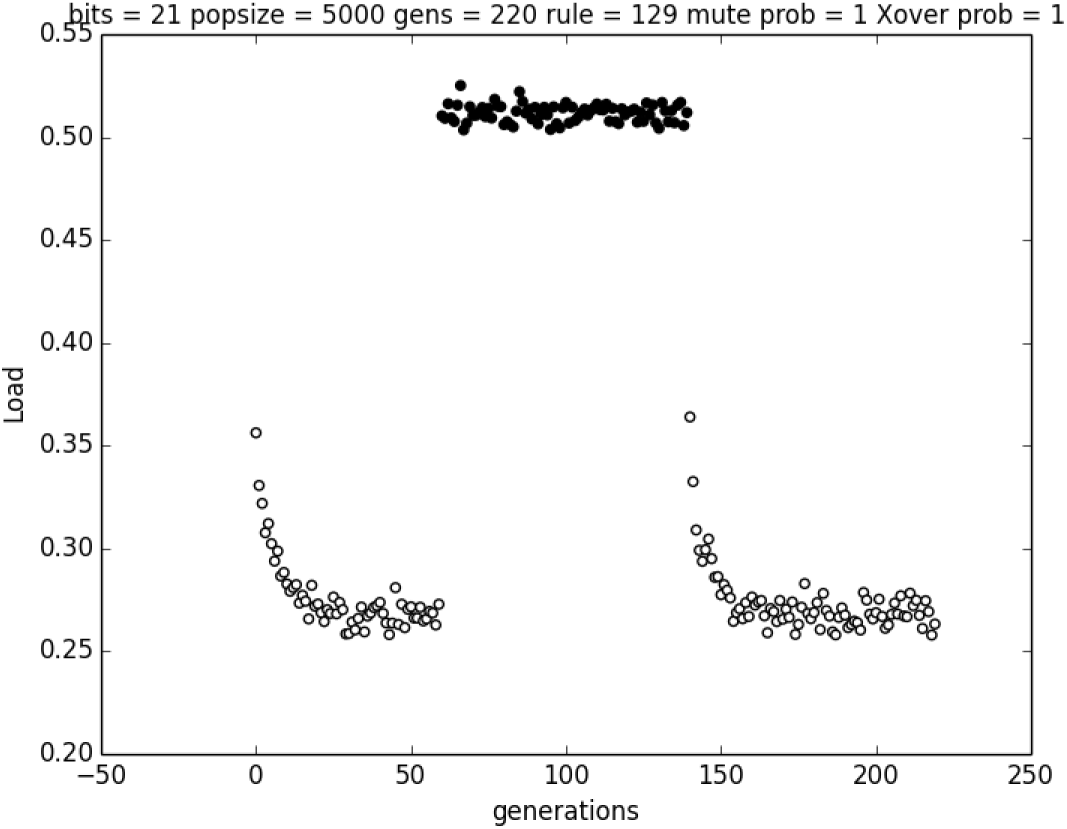
Rule 129 at 21 bits.This rule responds negatively with an increase in mutational load following removal of recombination

### 4.2 Survey of Networks

The 13 bit networks were generated for every rule that accepted more than zero genomes. Broadly speaking the graphs could be easily classified Fig. 10. Machines that generated one regular graph, multiple regular graphs but of the same order, regular graphs of differing order, irregular graphs with fur, through to connected furry graphs. Regular graphs occur when a rule accepts genomes with completely neutral regions. If a genome has *n* neutral bits then it will generate a regular graph with 2*^n^* nodes with n degree for each node. Rule 60 generates multiple regular graphs with the same degree. This shows that rule there are the same number of neutral bits in each regular graph but they are a different set of bits in each regular graph.

**Figure 10:**
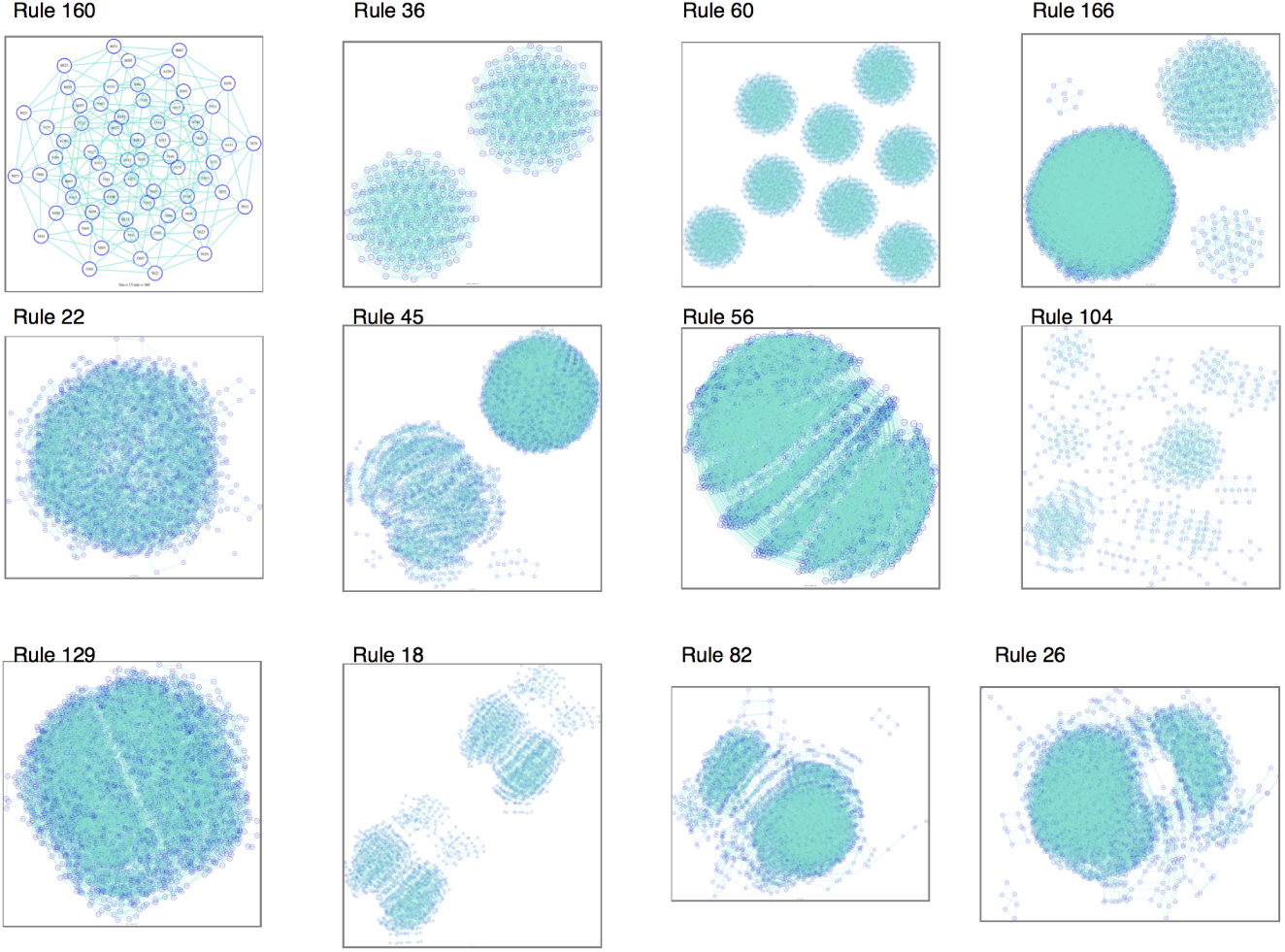
Sample of some of the 13 bit neutral networks produced, in order of increasing complexity

Rule 166 shows regular graphs of differing degree. The NofN slope for this graph is 1.0, this graph responds very quickly to mutation but had very little response to recombinational load. As the graphs become more complex they appear to respond better to recombination an exception being rule 129.

The results of the survey of the networks is shown in Fig. 11. Slope and S.E are correlated as the slope approaches 1.0 the variation decreases and as the slope approaches 0 the variation increases. Rules that generated only regular graphs of the same size have no variation in NofN so are not investigated any further by this means. Rules generating regular graphs of differing sizes with no fur or bridges give a slope of 1.0 as all connected nodes have the same number of neighbours. None of the NofN graphs produced in the survey appeared similar to that of sha256 however the similarity in the response to recombination between sha256 and some of the rules suggests that the rule set did contain class III machines. Indeed rule 30 at longer bit lengths is considered to be a PRNG.

**Figure 11:**
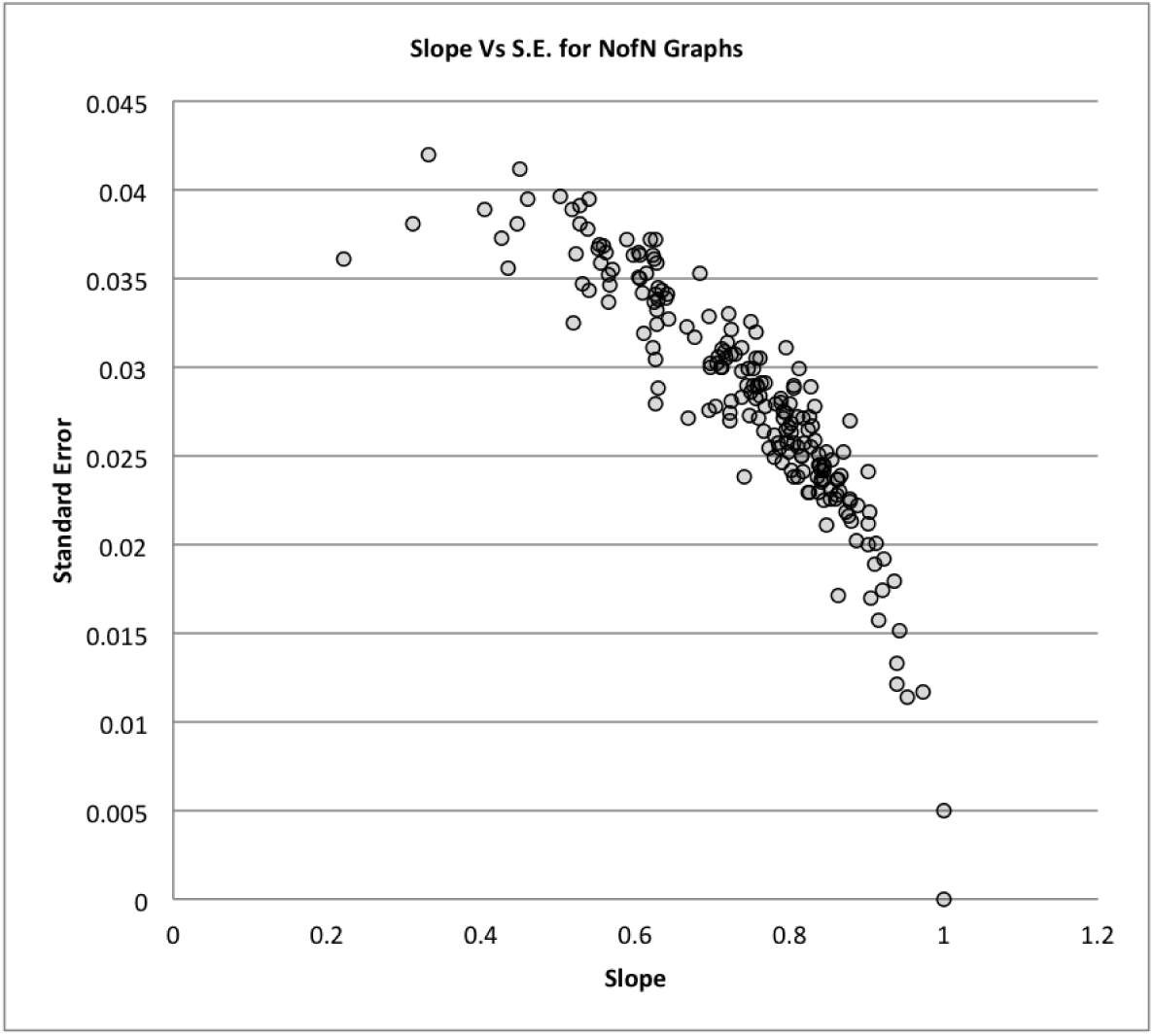
Slope vs Standard Error of the slope

### 4.3 Recombinational Centring RC

The phenomenon where recombination selects genomes that are more resistant to mutation than those produced through selection for reduced mutational load has been observed previously in model proteins(17) and gene artificial gene networks(25). It is not clear why RC occurs, it is likely due to the high degree of recombinational load experienced by poorly connected nodes in genome space. Recombinational load is likely to be a function of the hamming distance between two genomes, recombination rate and the density of solutions close to either parent. If there is one central mass of well connected nodes then the hamming distance between well connected genomes is rather small and the density of solutions is high, whereas the hamming distance between two randomly selected genomes in low density regions is likely to be high. The recombinational load for well connected nodes is much smaller than for poorly connected nodes and the viable recombinant offspring of a well connected node mated with a poorly connected node is more likely to be found close to the well connected parent. Recombination preferentially eliminates poorly connected nodes and moves the population into the denser regions of genome space for a given rule.

Those graphs that responded well to recombination did not all respond at the same rate. Rule 84 took approximately 10 generations to reach equilibrium where as rule 110 took nearly 80 generations to reach equilibrium. For some rules the drop in mutational load after recombination was short lived in others it was longer lasting.

### 4.4 Resistance to Ageing

Rules responded to ageing in different ways. Most rules saw and increase in resistance to ageing under asexual selection compared to the random starting population, however there was more variation in the way rules responded to sexual reproduction compared to asexual reproduction. Some rules greatly improved their resistance to ageing under sexual reproduction whereas at the other extreme sexual reproduction reduced resistance to the same as the random start population Fig 12.

**Figure 12:**
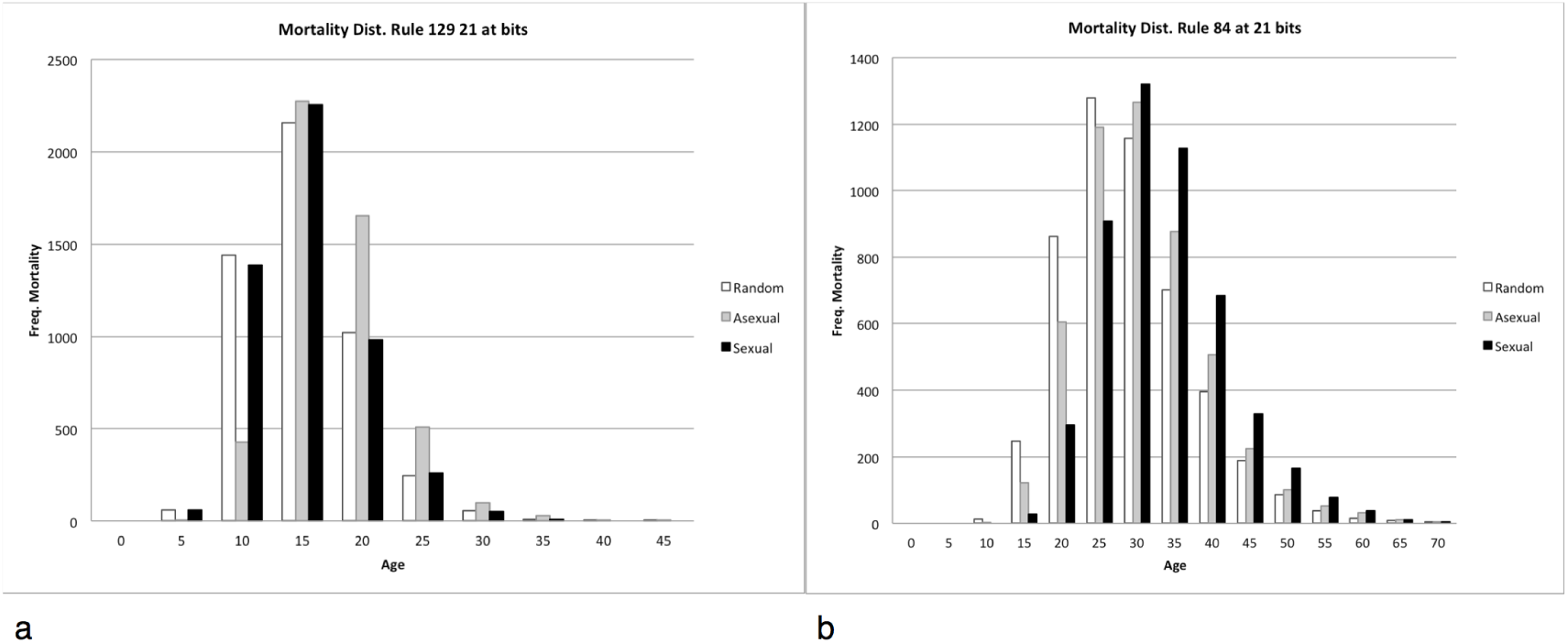
Response to ageing. a) shows a negative response to sex typical of a class III PRNGs. b) shows a dramatic positive in resistance to ageing seen in class IV rules

Resistance to ageing was measured by comparing the life span of the population at the end of 60 rounds of asexual reproduction with the lifespan after 80 rounds of sexual reproduction. The means of the two were compared and the result divided by the standard deviation of the sexual population

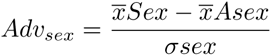

If sex is advantageous for the rule and the parameters used then *Adv*_sex_ will be greater than zero if sex is not beneificial then *Adv*_sex_ will be negative

The response to recombination in terms of longevity may correlate with the slope of NofN graph. There appears to be a weak relationship between the standard error of the slope and *Adv*_sex_ however values of SE that supported a good response to recombination were close to those that responded poorly Fig. 13. This is not surprising as class IV machines are on the edge of chaos and inevitable close to class III machines that are chaotic.

**Figure 13:**
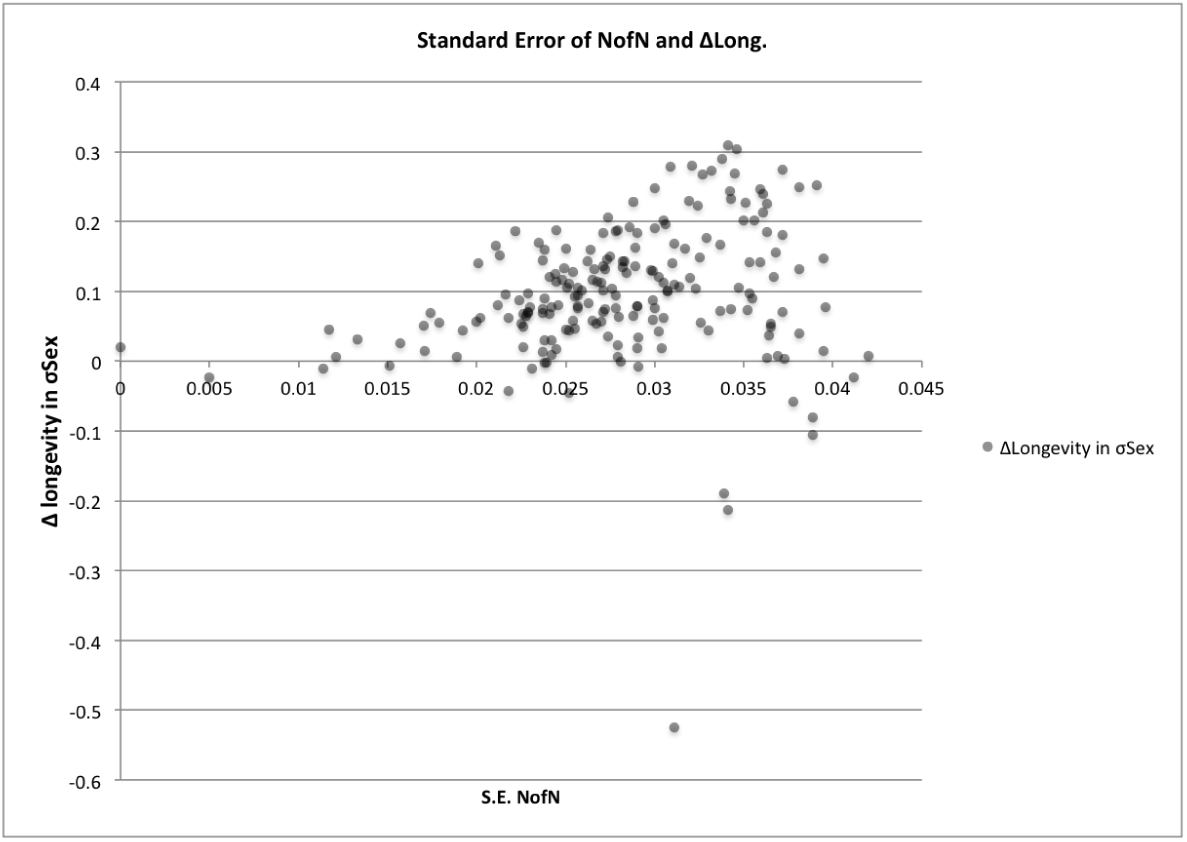
Standard errors relationship to the *Adv*_sex_ peaks at around 0.035 near the edge of chaos, close to rules giving a negative response to recombination suspected of being PRNG

### 4.5 Langtons Lambda

The onset of chaotic behaviour is often observed in complex systems however it is not easy to determine under what circumstances it occurs. Various parameters have been proposed to indicate which rule sets should give us edge of chaos behaviour. One that is easy to calculate but gives only a crude approximation.

Langtons *λ* parameter can be calculated thus.

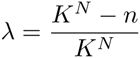

Where *n* is the actual number of the possible neighbour states used in the transition function. If *λ* was greater than 0.5 then 1 − *λ* was used for *λ* as the black white equivalent rule has the same behaviour but a lower *λ* value. While it is not possible to say with certainty which of the wolfram classes a given ECA rule belongs too. Langtons *λ* parameter gives some indication of the complexity of the machine used to accept the string (26). As *λ* approaches 0.5 the response to recombination becomes more positive with increasing complexity with *λ* =3*/*5 having the best response to sex Fig. 14. Rules with a negative response to recombination were found in 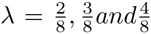. However rule 30 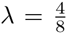 which is known to be a PRNG does not show PRNG behaviour at small genome lengths. At a genome length of 51 bits PRNG behaviour is found and a negative response is observed. So some of the rules are likely to respond more negatively at longer genome lengths

**Figure 14:**
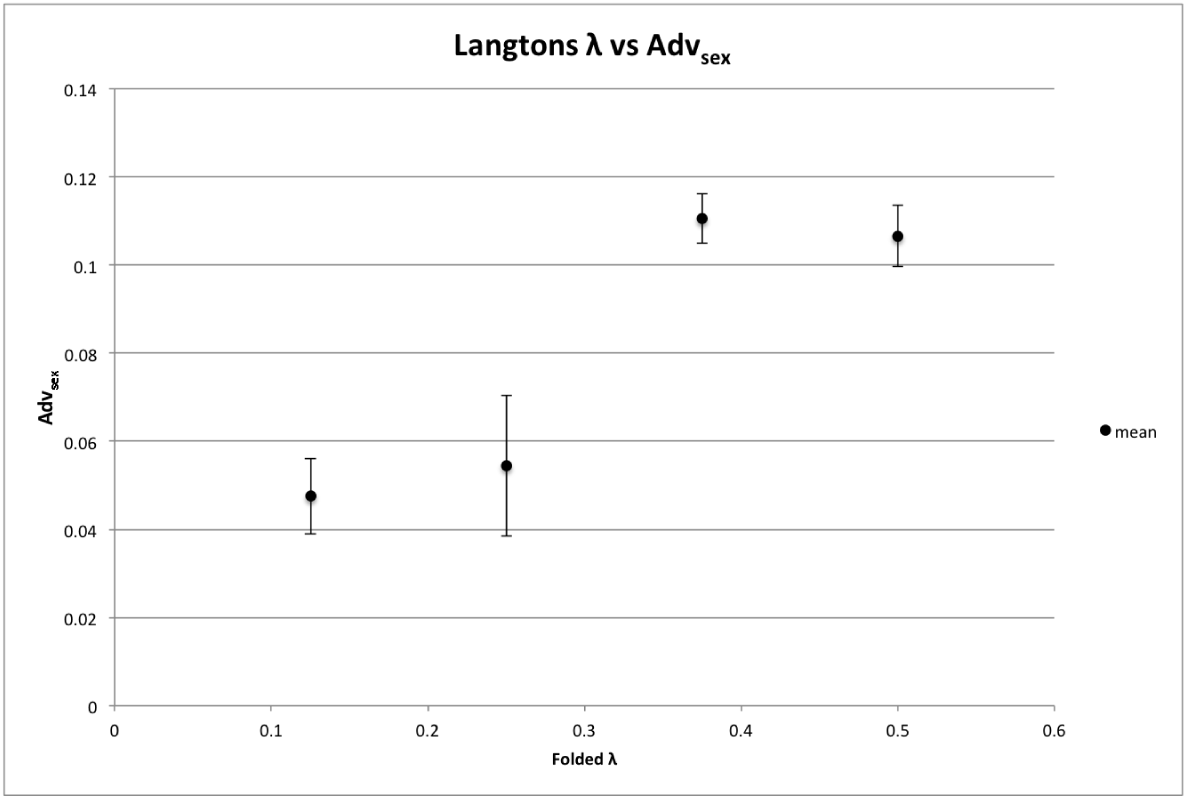
Rules with Langtons 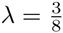 appear to be most likely to respond to recombination

### 4.6 Activity Time Portrait

In order to try and elucidate why some machines were responding to recombination and others weren’t, the behaviour of the compression process was investigated. This was achieved by superimposing all all the compression steps of the machine for all the members of a population at a given stage of the breeding system Fig. 15. The start population in the example shown is random showing mostly grey at the first line of the machines compression. Some pattern can be shown further down with the last bit on the bottom is of course always black as only viable individuals are used in the start population. Asexual reproduction sees some pattern appearing in the first line with some pattern developing in the processes towards the single last line single bit. After sexual reproduction the pattern comes more into focus with a clear pattern emerging. Some of the first line bits are completely white and some are completely black. This suggests the population is closely linked to a central archetype genome that has a high probability of viability. Relaxation of the pattern is seen when sexual reproduction is removed with the contrast between bits dropping down to asexual reproduction pattern.

**Figure 15:**
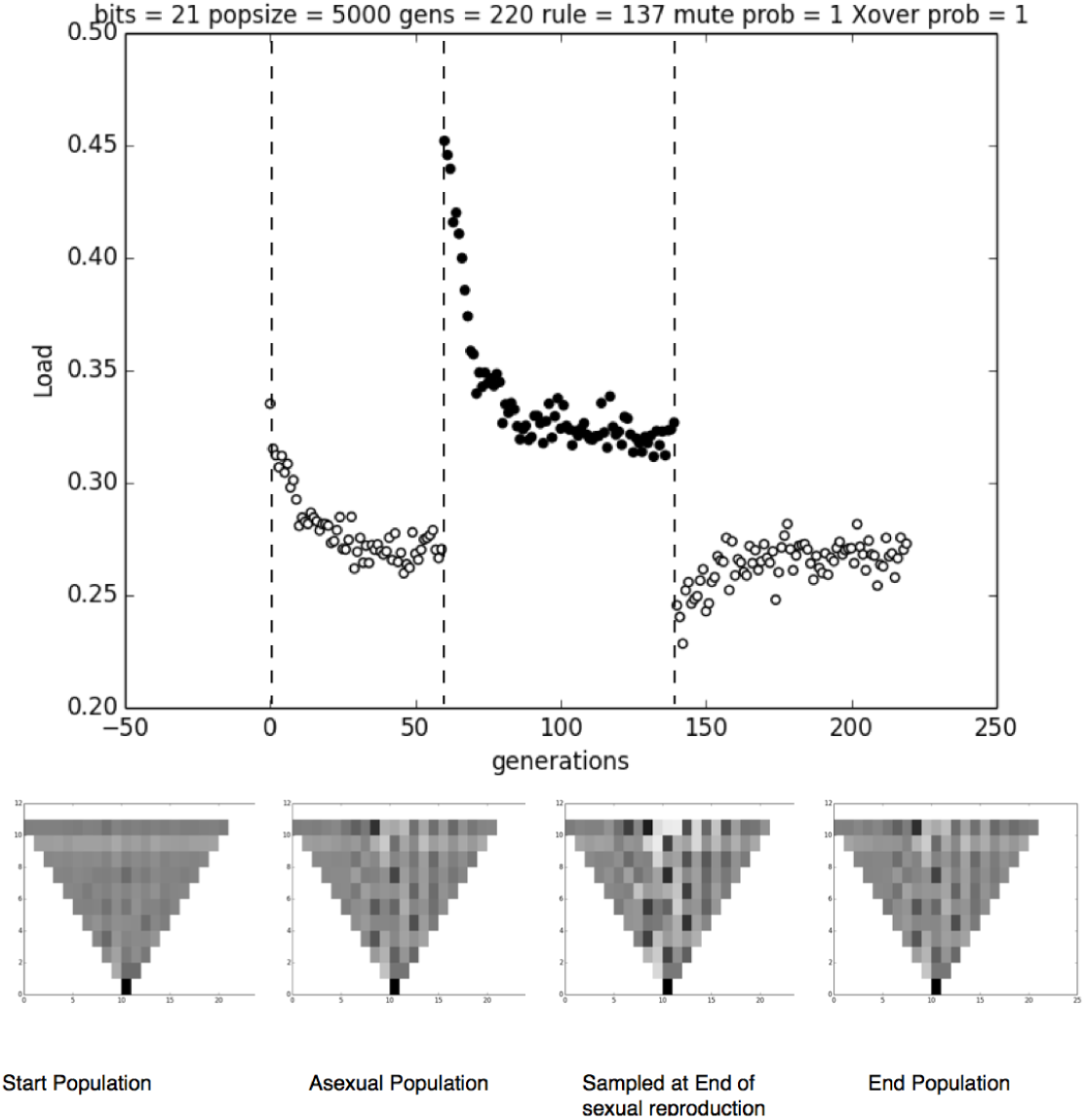
The activity portrait for the starting population, at asexual equilibrium, sexual equilibrium, and the end population

**Figure 16:**
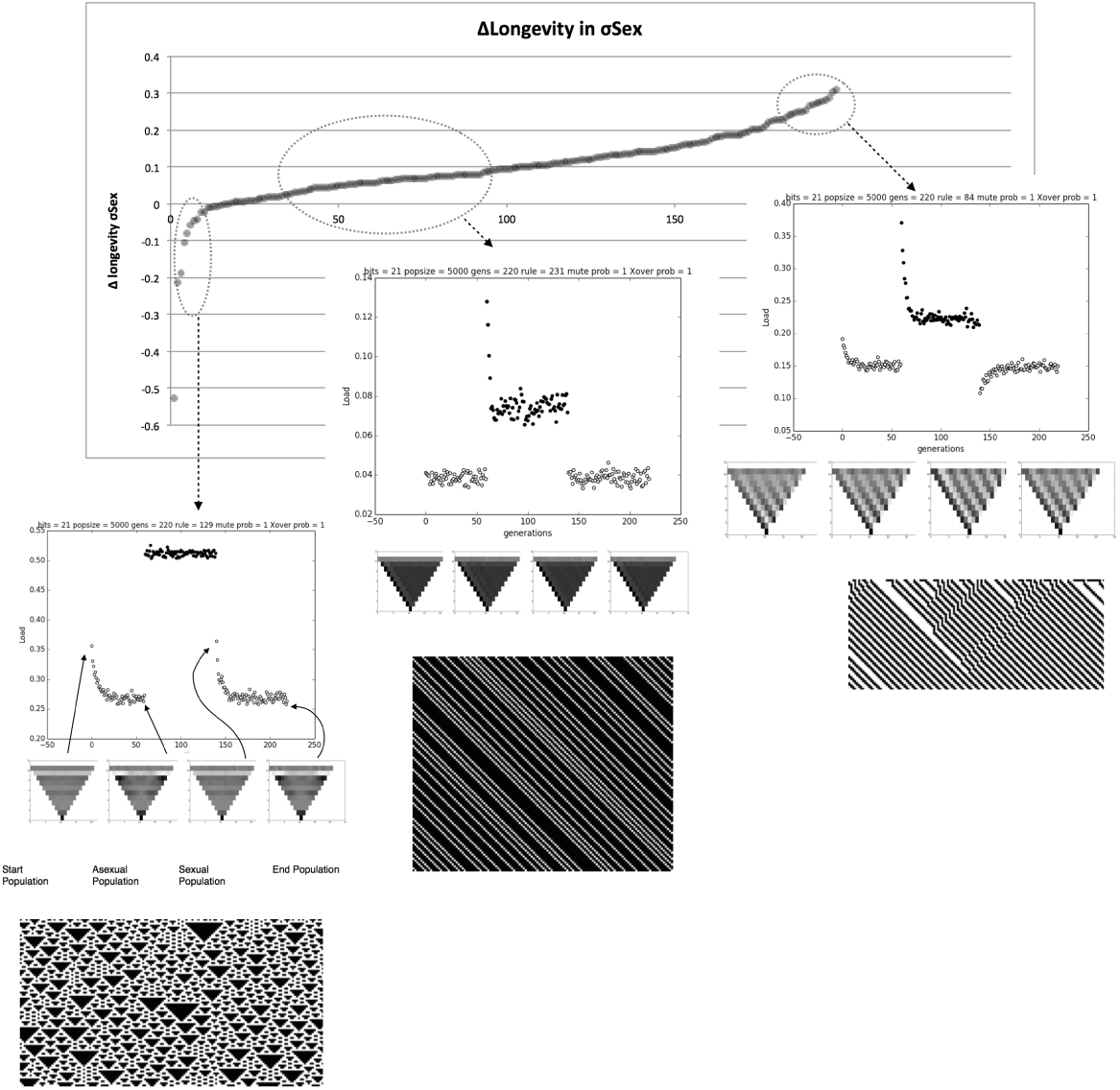
Each rule ranked by *Adv*_sex_ showing the activity portraits for the starting population, at asexual equilibrium, sexual equilibrium, and the end population at either extreme of the *Adv*_sex_

### 4.7 Activity Time Portraits and Advantage of Sex

Activity time portraits of every machine were made at the same stages of the breeding program. It was noticed that the machines that responded well to recombination in terms of longevity had more structure in the compression steps and had better contrast in the ATPs after recombination. In machines that responded negatively to sex the opposite was true. The first line had no, or weak patterns at every stage of the breeding program and recombination destroyed what pattern there was. This showed a striking similarity to what was seen using sha(256) kryptographic has function. Sha(256) is a very complex but completely deterministic machine, suggesting that class III machines that generate chaotic behaviour and are close to being pseudo random number generators respond badly to recombination. Class IV machines that have interacting gliders should show patterns seen in rule 137. The presence of well defined patterns in the machines with greater longevity due to recombination, suggests that gliders present in the genome are gliding down through the layers of compression in the nested machine and setting the last bit.

Recombination acts to reinforce these gliders by removing negatively interacting gliders and adding reinforcing gliders giving greater pattern in the ATP.

### 4.8 Intermittent Sex

The breeding experiments carried out used eighty cycles of sexual reproduction to show the drop in mutational load following recombination. Multicellular organisms of course do not undergo sexual reproduction every cell division. Some have a sequestered germline while others have a branching growth habit wilt multiple sexual organs per individual. Mutation accumulation must occur in the growth of the germ line. To investigate weather the recombinational centring process could tolerate mutation accumulation through vegetative growth, experiments were performed with intermittent sex. Here sex occurred every fifth generation instead of every generation Fig. 17. It was observed that as long as the period of vegetative growth was less than the time taken for the population to return to mutation alone levels of load, intermittent sex was able to lower mutational load and increase resistance to ageing

**Figure 17:**
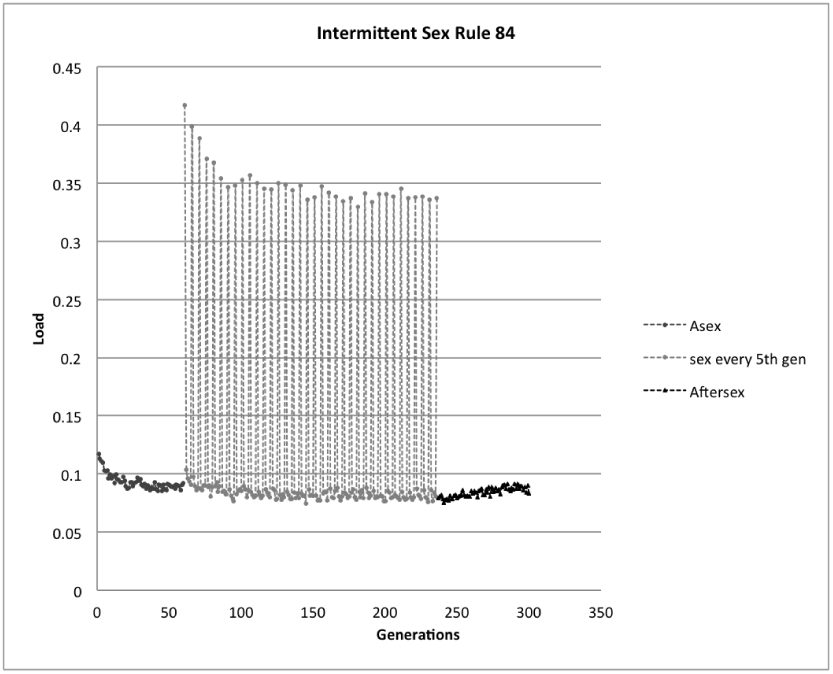
Sex every fifth generation is sufficient to condense the population into the dense region of genome space.

## 5 Discussion

The evolution of sex is an ongoing mystery in science. Maynard-Smith’s insight into the two fold cost of sex and question of why sexual populations are not invaded by asexual mutants set the stage for attempts to answer this question. Most theories have attempted to answer this question using traditional population genetics where individuals are treated as single diploid genomes with a small number of loci. In contrast the model presented here uses the whole genome and considers the life history of an individual.

With the exception of Khondrashov’s deterministic mutation hypothesis all the theories require extrinsic factors such as environmental variation and parasitic interactions to explain maintenance of sex. These theories have some overlap in their ideas however they all require genes and fitness to be real entities. Fitness is not a real measurable property of individuals as has been discussed by Richard Dawkins in chapter 10 of The Extended Phenotype (27). It can only be applied to features of individuals that may change the probability of survival. For example, Individuals with red wings may survive more often than individuals with green wings but that does not tell you if a green winged individual is going to survive or not because that depends on the biological background in which those wings are found.

Here it is proposed that recombination increases longevity in complex organisms and this is sufficient to compensate for the costs associated with sexual reproduction. This theory relies on the intrinsic factors of an individuals growth, development and senescence.

A multicellular organisms first problem is to complete development and then to resist entropy as long as possible (28) so as to reproduce. All through that process genetic and epigenetic information is being inherited from cell to cell via cell division with some error. Multicellular organisms are not a point in genome space but a cloud of points with a scale of mutations running the spectra from the nucleotide level, up through, duplications, deletions, translocations and inversions, to chromosomal abberatons. These changes can be neutral, or result in hypoplasia, hyperplasia, or inappropriate expression of cell products. Shared fate is the fundamental problem of multicellularity (9). Each cell is dependent on other cells for its survival, defection of one cell can kill all the cells. This pressure is always present on every individual in every species of multicellular organism.

Recombination has already been shown to increase thermodynamic stability in protein like models (17). Proteins are made up of interacting residues that define the structure. This concept of interacting components is found at every level of biological complexity, protein domains have to interact appropriately as do proteins, cells, tissues and organs. Cell and tissue specialisation requires that there be synergy between these components and variation in these components will have complex effects. However mutation can simulate environmental perturbation at higher levels just as mutation can simulate thermodynamic instability at the molecular level. Resistance to perturbation requires the presence of many alternate pathways and redundant systems, producing high levels of conditional neutrality, where mutations are only be exposed to selection in the presence of other mutations. One of the criticism of the Deterministic Mutation Hypothesis is that the mutation rate observed in nature is not high enough, however this can be countered by the somatic mutation rate which must also be considered and the increase in robustness independent of the probability of mutation.

The wolfram ECAs have been modified in this experiment to generate grammars that accept or reject strings based on the interactions between various bits of the string governed by the rule. Wolfram and others have proposed that cellular automatas fall into four categories. Culik (22) has shown that placing each ECA into a particular class is undecidable. This does not mean that we can’t classify come ECAs it means there is no algorithm that will place every ECA in one of the four classes. As pointed out by Langton (26) one of the main insights of the Wolfram classes and ECAs is that there is an upper bound on complexity, the pseudo random number generator or PRNG. This idea can be applied directly to biology particularly with regard to epistasis. The concept of positive and negative epistasis has been developed over the years to cope with selection on multiple genes. However it cannot be applied to individuals as the idea requires fitness to be a real property of genomes. This breaks down once genome size reaches any appreciable size because genome space expands exponentially with genome length. At any reasonable genome length, genome space reaches cosmic proportions and any practical population of individuals becomes minute in comparison. It is extremely unlikely to have two individuals with the same genome even in single celled organisms, therefore we can never know fitness of an individual genome in terms of probability of survival.

The class III and class IV machines however are good ways of looking at epistasis in genomes. In class III machines that are either PRNGs, or close to them, every mutation has a 50 percent chance of mortality. These machines are critically built and are uniquely unsuitable for use in multicellular machines, as offspring of a cell will have highly variable viability and low resilience. Class IV machines can undergo recombinational centring thus increasing robustness and are more resilient to time dependent intrinsic factors such as accumulation of genetic and epigenetic errors and well as environmental perturbation. There are however some anomalous results regarding rule 110 Fig. 18. It was expected that gliders would increase the activity of recombinational centring as long as the glider is able to set the final bit and its length is short relative to the genome length. However rule 110 which has a plethora of gliders and glider interactions has an unusual response to recombinational load. There is an alphabet of gliders and they have a neighbourhood of gliders they can interact with, effectively building another machine at a higher level. As we know that rule 110 is a universal machine (29) and can therefore simulate any other rule including PRNGs.

**Figure 18:**
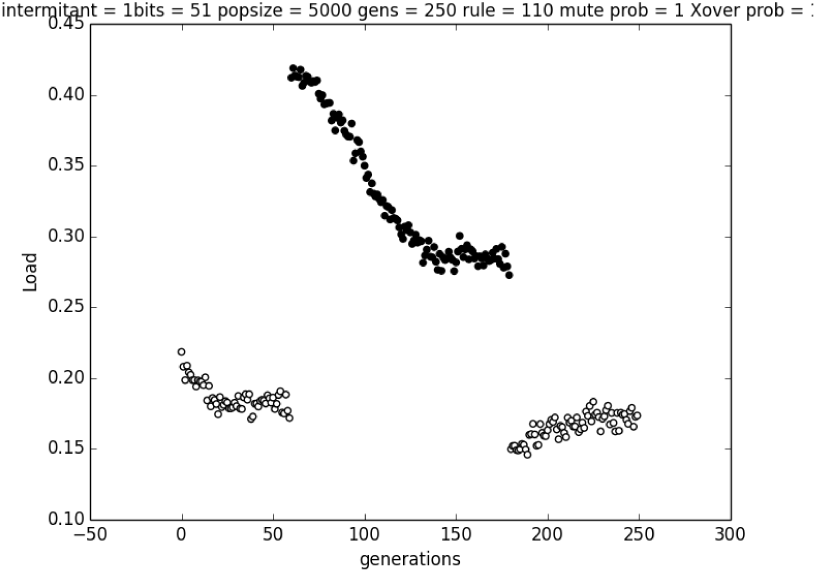
Rule 110 at 51 bits, It takes a long time to reach equilibrium under recombinational load. The process seems to have several phases and under asexual reproduction takes quite some time to return to asexual equilibrium

### 5.1 Genes as Gliders

Genes, in a information sense, are simply substrings of strings. Logically substrings can also have substrings and this is what we see in molecular biology with genes having domains, motifs etc. It is the biological machinery that confers discrete boundaries to gene sequences. Just as proteins are built up of amino acids in a sequence specified by the DNA of a cell so gliders are built up of bits of the genome of the machine. We can therefore view genes as gliders, they are generated by the machine and interact with each other to decide phenotype and ultimately decide viability. There may be two types of gliders for a given rule, positive gliders that will glide down and set the viability bit to 1 and negative gliders that will set the bit to 0. Negative gliders are of course lethal. If a genome is being made viable by the action of positive gliders then as genome length increases so does the number of neutral mutations as only the glider is setting the viability bit. Gliders may interact in a variety of ways. They may reinforce or annihilate each other. Rule 110 has large range of gliders and glider interactions (30). If biological machines are at least as complex as rule 110 then biological systems are universal computers and are thus equivalent to turing machines. The relationship between genome and phenotype must therefore be undecidable, that is we cannot generate any algorithm that when given any machine and a genome it will tell us its phenotype. This does not mean we cannot calculate the phenotype of some of the genomes, just as we can program computers despite the halting problem being undecidable, it is just not possible to write a program that will tell you phenotype of every genome. The other consequence of being a turing machine is that for most inputs, the machine behaves as a PRNG, this may explain the unusually long time it takes for rule 110 to reach equilibrium under recombination.

### 5.2 Inbreeding Depression

Although this system is haploid it may show some light on the behaviour of diploid systems, although care must be taken to predict how this systems would behave if diploidy was added to its design. Inbreeding depression is a well known phenomenon of diploid organisms and is usually considered to be due to the loss of heterozygosity and the benefits of heterosis. However the drop in vigour observed in this system shows that even haploid systems may show inbreeding depression. Experiments on inbreeding necessarily require selfing or sibling mating to remove heterozygosity. Reduction in effective population size results in differences in genetic load becoming nearly neutral. The population can now roam randomly through genome space resulting in a the decrease in longevity and fecundity which is often observed (31) and in some cases changes in resilience to stress (32).

### 5.3 Benefit of Longevity

Can increased longevity compensate for the costs of sex. If maturation time is long enough then presumably at some point the reproductive excess of sexual versus asexual individuals may be more than two fold. The difference in resistance to ageing will result in a greater reproductive potential which can be approximated as the area under the mortality curve past the maturation age Fig. 19a.

**Figure 19:**
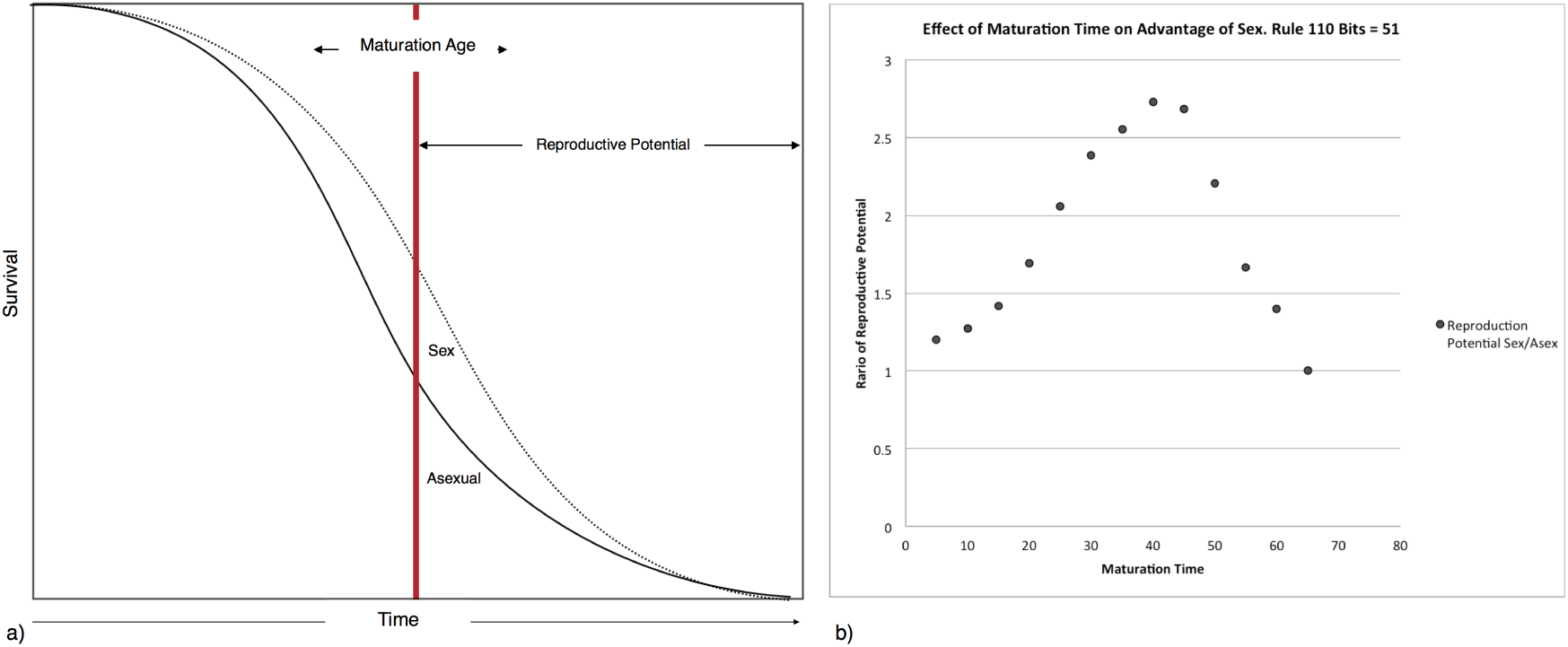
a) Schematic showing the difference in the area under the mortality curve of the sexual and asexual populations for a given maturation age. b) The affect of maturation age on the ratio of the area of the curves above maturation age for sexual and asexual population for rule 110 at 51 bits

When the ratio of areas under the curve for various maturation ages was compared for rule 110, maturation times from 40 to 50 units gave advantages greater than 2.5 times asexual reproduction for sexual reproduction Fig. 19b. This suggests that sexual reproduction may, under the right circumstances, be able to compensate for the two fold cost of sex by increased reproductive potential in long lived organisms through extension of reproductive life span.

The recombinational load aspect may be more complex, however in most of the machines showing a benefit of recombination the recombinational load eventually fell to quite low levels. However there is usually a difference between the two loads. The cost of recombinational load may be paid at the gametic stage or at the early embryo stage, indeed there is a correlation between chiasma frequency and log gamete redundancy (33) with an exponential increase in gametes produced per fertilisation compared to increasing chiasma. Most large animals produce a vast excess of sperm and the effect of recombinational load on fertility would be insignificant. There is variation in the quality of sperm produced, however, it is not clear if this is due to variation in the haploid genome.

There is also potential for genetic load to be paid at the embryo stage. Even in humans, which historically could have more the ten pregnancies in a life time, two thirds of embryos fail to reach live births (34), mostly due to chromosome abnormalities that occur after fertilisation (35). The very high level of early embryo mortality in humans is quite surprising and appears to be due to the instability of cell division early in embryogenesis, but it indicates that, even in animals with small litters, a high degree of prenatal mortality can be tolerated.

### 5.4 Epistasis and the Edge of Chaos

The role of epistasis in evolution has been well covered in the literature (36). Negative epistasis is used as an argument for the deterministic mutation hypothesis (DM)of Kondrashov. Negative epistasis is where the effect of two mutations on fitness is greater than they are individually, positive epistasis is where the effects of mutations are less severe than expected. It is challenging to map the epistasis concepts of negative and positive epistasis to the nested rule systems in this paper and wolfram classes of ECA. This is partly due to the confusion produced using the imaginary concept of fitness as a real property of individuals and the model of evolution as change in allele frequencies. In this paper “fitness” is binary with only viable and non-viable forms and evolution is seen as a journey through genome space with selection acting to move the population to denser regions of genome space. The density is proportional to the redundancy, which is equal to the robustness or stability of that region of the genome space. Redundancy cannot be selected for at the “gene” level as the coefficient of selection is dependent on the state of a genome at another locus. The DM hypothesis requires negative epistasis, where sublethal mutations are lethal in the presence of other lethal mutations. I show that recombination searches genome space for high density regions and can discover high density regions as has also been seen is searches of protein sequence space (17).

Consider two regions of genome space each with a glider at a different locus. One glider will be called “Belt” the other “Braces” Fig. 20. Genomes with either “Belt” or “Braces” are viable as mutations have to hit and disable either of the gliders, so most mutations are neutral and thus there is a high density of solutions and low mutational load. Recombining the two gliders results in a genome with belt and braces, if these do not negatively interact then mutation of either of these individually is neutral. At least two mutations are required before lethality can be achieved. In addition this region of genome space has low recombinational load as both belt and braces would have to be swapped out for lethality to occur. In addition to redundancy through positive interactions redundancy could also be achieved through mutation suppression. A mutation that generates a glider that annihilates negative gliders, that is gliders that will flip the viability bit to zero, will render those negative glider mutations neutral.

**Figure 20:**
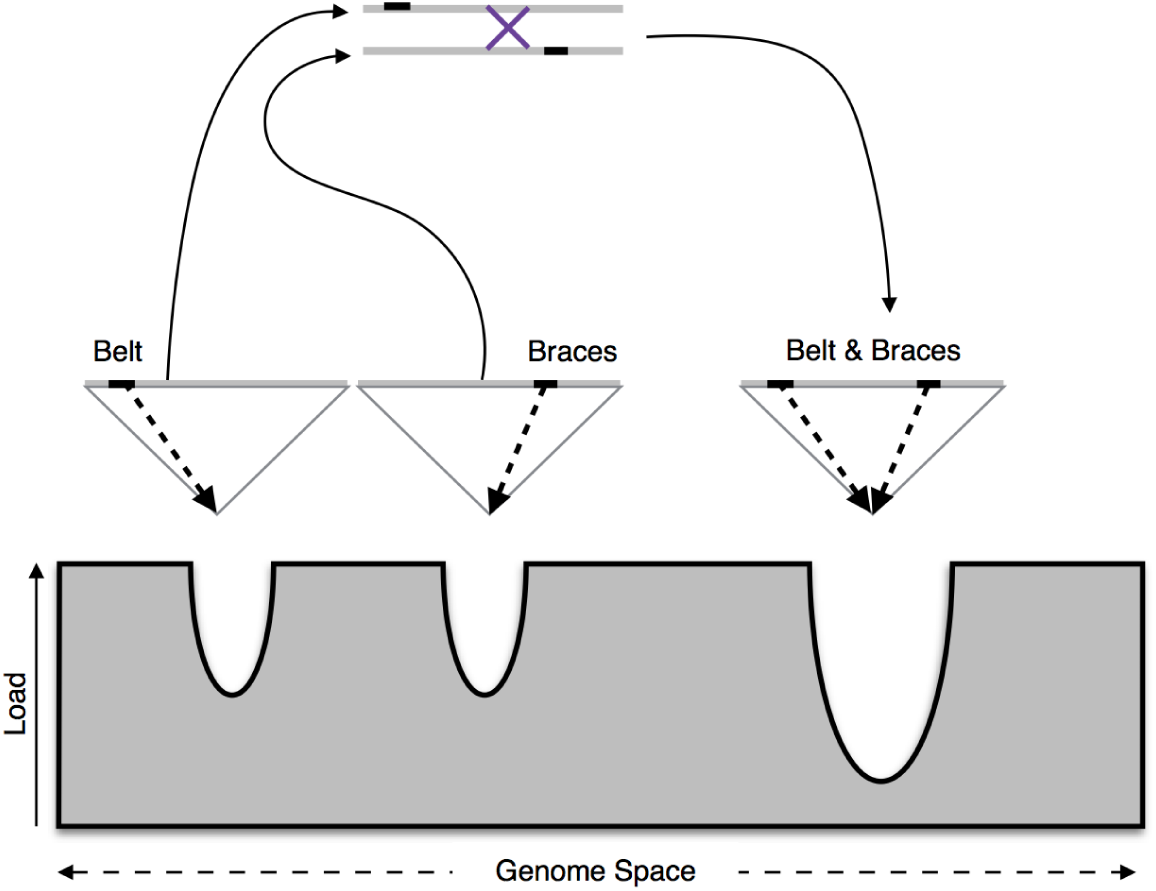
A genome with a “Belt” glider and a genome with a “Braces” can recombine to form a genome with “Belt” and “Braces” which has a greater density and lower load

There must be some relationship between epistasis and ageing. Epistasis curves show the response to mutation accumulation. If there is high redundancy then it will take time for the accumulation of mutations of have an affect, this is negative epistasis. Positive epistasis is some what similar to a PRNG where mutations have large effects early on in ageing. It therefore seems likely that all large complex multi-cellular organisms require negative epistasis.

Returning to the classical models of epistasis used in the theory of the evolution of recombination, we can see that recombination seeks to make epistasis more negative leading to increased resistance to ageing.

Class IV machines generate neutral networks with correlated mutational load, this make multi-cellularity possible. Class III machines generate poorly correlated networks, multi-cellularity is severely limited in such machines as they can only decrease their load through mutation as sex has a negative effect on viability. Class III machines would there for be limited to small, asexual multi or single celled organisms.

### 5.5 Ageing in Large Multicellular Organisms

It appears that ageing rates and life history are correlated according to Ricklefs (37). Mammals with longer gestation periods having slower ageing rates and with both birds and mammals showing an inverse relationship between ageing and gestation or incubation period, and age of maturity. It has been pointed out by Burt and Bell (38) that there is a correlation, at least in mammals, between the amount of excessive chiasma and age at maturity in non-domesticated species, a similar pattern has been observed in marsupials (39). Increased recombination rates in larger organisms would result in greater concentration of the population in the dense regions of genome space. This would result in greater resilience and a higher probability of completing development and increased longevity.

Large organisms are presumably harder to make as more cell divisions need to occur in order for development to complete. Larger size increases the opportunity for accumulation of error, these errors are unpredictable at the small scale but predictable in their ultimate consequence. large complex organisms composed of integrated machines and myriads of cells and genome copies, ultimately face an error catastrophe due the process of growth through cell proliferation. Large organisms, necessarily also have more cell divisions in their germline, however as show in the intermittent sex model, as long as the germline cell divisions are insufficient for a return to the asexual level of load, improvement in resistance to ageing can still occur through periodic recombination.

Ageing in humans follows the Gomepertz Markham law, where the probability of death is log linear over a certain age. This gives a doubling of the probability of death approximately every 8 years. It has been suggested that the Gompertz curve is a special case of the Weibull curve based on reliability theory (40) (41). Gavrilov and Gavirlova have proposed that the ageing pattern, can bee seen as the consumption of redundancy over time in a population with varying levels of intrinsic reliability.

Both drosophila and humans share the same “bath tub” curve of mortality where there is high levels of failure early on, that drops down in the early stages of life and then increases with the typical Gompertz curve (40), this is even more evident if prenatal mortality is considered in humans. In reliability systems this is seen as the “working in” or “burning out” of defective components. In biological systems this can probably due to two causes, recombinational load and mutational load combined with the sensitivity of development to errors in the developmental process. It has recently been shown, in Drosophila, that individuals with greater developmental symmetry also have greater longevity (42) this is consistent with this reliability model where reliability early on is conferred by the same factors providing resistance to error in later life.

Cancer turmourigenesis is a multistep process requiring the acquisition of several traits (16). There are layers of resistance to cancer growth each of these must be overcome for tumour growth to continue. Individuals bprn with some of the barriers missing are not only more likely to develop cancer but are less likely to complete development correctly. The ability of recombination to concentrate a population in the highly redundant centre of neutral network is consistent with the view of reliability theory and ageing proposed by Gavrilov and Gavrilova.

The timely and accurate completion of development is the first problem faced by a multi-cellular organism. From the first cell division, entropy is increasing in the individual as the errors of cell division are added to those inherited from either parent. Resistance to recombination selects regions of genome space with high redundancy, this results in organisms with highly robust genomes, more likely to complete development and to be long lived. The increase in longevity may compensate for the costs of sex, including the two fold cost of sex, through increased reproductive potential in long lived organisms requiring long maturation times. Large complex species are therefore resistant to invasion by asexual mutants, whereas small simple organisms with early maturation should be vulnerable to invasion by asexual forms.

## 6 Acknowledgements

I would like to thank Bakhadyr Khoussainov from the computer science department of the University of Auckland for his help in developing some of the ideas presented in this paper.

## References

[1] G. Bell, The masterpiece of nature: the evolution and genetics of sexuality. CUP Archive, 1982.

[2] J. Maynard-Smith, The evolution of sex. Cambridge Univ Press, 1978.

[3] R. A. Fisher, The genetical theory of natural selection: a complete variorum edition. Oxford University Press, 1930.

[4] H. J. Muller, “Some genetic aspects of sex,” American Naturalist, pp. 118–138, 1932.

[5] S. P. Otto and N. H. Barton, “Selection for recombination in small populations,” Evolution, vol. 55, no. 10, pp. 1921–1931, 2001.

[6] A. S. Kondrashov, “Deleterious mutations and the evolution of sexual reproduction,” Nature, vol. 336, no. 6198, pp. 435–440, 1988.

[7] S. P. Otto and A. C. Gerstein, “Why have sex? The population genetics of sex and recombination,” Biochem. Soc. Trans., vol. 34, no. Pt 4, pp. 519–522, Aug 2006.

[8] E. Szathmary and J. M. Smith, “The major evolutionary transitions,” Nature, vol. 374, no. 6519, pp. 227–232, Mar 1995.

[9] L. W. Buss, The evolution of individuality. Princeton University Press, 1987.

[10] H. A. Orr, “Somatic mutation favors the evolution of diploidy,” Genetics, vol. 139, no. 3, pp. 1441–1447, Mar 1995.

[11] J. W. Drake, B. Charlesworth, D. Charlesworth, and J. F. Crow, “Rates of spontaneous mutation,” Genetics, vol. 148, no. 4, pp. 1667–1686, 1998.

[12] D. R. Denver, K. Morris, M. Lynch, and W. K. Thomas, “High mutation rate and predominance of insertions in the caenorhabditis elegans nuclear genome,” Nature, vol. 430, no. 7000, pp. 679–682, 2004.

[13] C. F. Baer, M. M. Miyamoto, and D. R. Denver, “Mutation rate variation in multicellular eukaryotes: causes and consequences,” Nature Reviews Genetics, vol. 8, no. 8, pp. 619–631, 2007.

[14] H. J. Muller, “The relation of recombination to mutational advance,” Mutation Research/Fundamental and Molecular Mechanisms of Mutagenesis, vol. 1, no. 1, pp. 2–9, 1964.

[15] C. Tomasetti and B. Vogelstein, “Variation in cancer risk among tissues can be explained by the number of stem cell divisions,” Science, vol. 347, no. 6217, pp. 78–81, 2015.

[16] D. Hanahan and R. A. Weinberg, “The hallmarks of cancer,” cell, vol. 100, no. 1, pp. 57–70, 2000.

[17] Y. Xia and M. Levitt, “Roles of mutation and recombination in the evolution of protein thermodynamics,” Proceedings of the National Academy of Sciences, vol. 99, no. 16, pp. 10 382–10 387, 2002. [Online]. Available: http://www.pnas.org/content/99/16/10382.abstract

[18] M. Eigen, “Viral quasispecies,” SCIENTIFIC AMERICAN-AMERICAN EDITION-, vol. 269, pp. 32–32, 1993.

[19] C. O. Wilke and C. Adami, “Evolution of mutational robustness,” Mutation Research/Fundamental and Molecular Mechanisms of Mutagenesis, vol. 522, no. 1, pp. 3–11, 2003.

[20] E. van Nimwegen and J. P. Crutchfield, “Metastable evolutionary dynamics crossing fitness barriers or escaping via neutral paths?” Bulletin of Mathematical Biology, vol. 62, pp. 799–848, 2000.

[21] S. A. Kauffman, The origins of order: Self organization and selection in evolution. Oxford university press, 1993.

[22] K. Culik II and S. Yu, “Undecidability of ca classification schemes,” Complex Systems, vol. 2, no. 2, pp. 177–190, 1988.

[23] S. Wolfram, “One-dimensional cellular automata,” 7 2015. [Online]. Available: http://atlas.wolfram.com/01/01/rulelist.html

[24] S. C. North, “Drawing graphs with neato,” NEATO User Manual, vol. 11, 2004.

[25] R. B. Azevedo, R. Lohaus, S. Srinivasan, K. K. Dang, and C. L. Burch, “Sexual reproduction selects for robustness and negative epistasis in artificial gene networks,” Nature, vol. 440, no. 7080, pp. 87–90, 2006.

[26] C. G. Langton, “Computation at the edge of chaos: Phase transitions and emergent computation,” Physica D, vol. 42, pp. 12–37, 1990.

[27] R. Dawkins, The extended phenotype: The long reach of the gene. Oxford University Press, 1999.

[28] L. Hayflick, “Entropy explains aging, genetic determinism explains longevity, and undefined terminology explains misunderstanding both,” PLoS genetics, vol. 3, no. 12, p. e220, 2007.

[29] M. Cook, “Universality in elementary cellular automata,” Complex Systems, vol. 15, no. 1, pp. 1–40, 2004.

[30] G. J. Martínez, H. V. McIntosh, and J. Mora, “Gliders in rule 110,” International Journal of Unconventional Computing, vol. 2, no. 1, p. 1, 2006.

[31] A. Hoikkala, M. Saarikettu, J. S. Kotiaho, and J. O. Liimatainen, “Age-related decrease in male reproductive success and song quality in drosophila montana,” Behavioral Ecology, vol. 19, no. 1, pp. 94–99, 2008.

[32] C. Vermeulen and R. Bijlsma, “Changes in mortality patterns and temperature dependence of lifespan in drosophila melanogaster caused by inbreeding,” Heredity, vol. 92, no. 4, pp. 275–281, 2004.

[33] J. Cohen, “Cross-overs, sperm redundancy and their close association,” Heredity, vol. 31, no. 3, pp. 408–413, 1973.

[34] N. S. Macklon, J. P. Geraedts, and B. C. Fauser, “Conception to ongoing pregnancy: the ‘black box’of early pregnancy loss,” Human Reproduction Update, vol. 8, no. 4, pp. 333–343, 2002.

[35] E. Vanneste, T. Voet, C. Le Caignec, M. Ampe, P. Konings, C. Melotte, S. Debrock, M. Amyere, M. Vikkula, F. Schuit et al., “Chromosome instability is common in human cleavage-stage embryos,” Nature medicine, vol. 15, no. 5, pp. 577–583, 2009.

[36] P. C. Phillips, “Epistasis—the essential role of gene interactions in the structure and evolution of genetic systems,” Nature Reviews Genetics, vol. 9, no. 11, pp. 855–867, 2008.

[37] R. E. Ricklefs, “Life-history connections to rates of aging in terrestrial vertebrates,” Proceedings of the National Academy of Sciences, vol. 107, no. 22, pp. 10 314–10 319, 2010.

[38] A. Burt and G. Bell, “Mammalian chiasma frequencies as a test of two theories of recombination,” Nature, vol. 326, no. 6115, pp. 803–805, 1987.

[39] P. Sharp and D. Hayman, “An examination of the role of chiasma frequency in the genetic system of marsupials,” Heredity, vol. 60, no. Pt 1, pp. 77–85, 1988.

[40] L. A. Gavrilov and N. S. Gavrilova, “Models of systems failure in aging,” Handbook of Models for Human Aging, pp. 45–68, 2006.

[41] T. B. Kirkwood, “Deciphering death: a commentary on gompertz (1825)‘on the nature of the function expressive of the law of human mortality, and on a new mode of determining the value of life contingencies’,” Philosophical Transactions of the Royal Society of London B: Biological Sciences, vol. 370, no. 1666, p. 20140379, 2015.

[42] L. G. Harshman, H.-G. M¨uller, X. Liu, Y. Wang, and J. R. Carey, “The symmetry of longevity,” The Journals of Gerontology Series A: Biological Sciences and Medical Sciences, vol. 60, no. 10, pp. 1233–1237, 2005.

